# Variable effects of biocontrol bacteria on potato resistance against black leg caused by soft rot Pectobacteriaceae in the field

**DOI:** 10.64898/2026.02.09.704818

**Authors:** Viola Kurm, Jack Gros, Carin Lombaers, Yvonne Griekspoor, Odette Mendes, Marjon Krijger, Patricia van der Zouwen, Leo Poleij, Jan van der Wolf

## Abstract

Blackleg disease in potato, caused by soft rot Pectobacteriaceae, is a substantial cause of loss in seed potato production. Recent research has attempted to identify bacteria with antagonistic activity against several diseases, among which blackleg. However, most biocontrol agents have been tested only *in-vitro* or in the greenhouse. In this study, we tested the effect of bacterial biocontrol agents in a four-year field experiment against blackleg caused by *Pectobacterium brasiliense* and *Dickeya solani*. Effects of the treatments on disease incidence was highly variable between years and also differed between cultivars, soil type and even replicates. Disease incidence was on average higher in sandy soil compared to clay soil and higher in the cultivar Kondor than Mozart. For a subset of the bacterial isolates genome mining could detect the presence of genes involved in the production of antibiotics and siderophores, but this was not correlated with disease incidence in the field. Moreover, most isolates were able to survive in storage on tubers from inoculation until planting. Thus, we conclude that while the used isolates showed the potential for antagonistic activity and were present on tubers when planting, no antagonist treatment could consistently decrease disease incidence. Inoculation of the isolates on the tuber surface might have been insufficient for plant colonization.

## Introduction

Soft rot Pectobacteriaceae (SRP) cause severe losses in international seed potato production as the causative agents of soft rot in storage and black leg in the field. In the Netherlands, the species *Pectobacterium brasiliense* and to a lesser extent *Dickeya solani* and *Pectobacterium parmentieri*, are the most frequently occurring SRP species (van der Wolf, et al., 2022). Although some potato cultivars are less susceptible to SRP, there are no resistant cultivars as of yet. Moreover, no chemical bactericides are known to be effective against this group of bacteria. Current mitigating measures include the use of sterile disease-free mini-tubers as an initial seed and appropriate storage conditions that prevent spread of the disease (Charkowski, 2015, Van Gijsegem, et al., 2021). However, disease caused by SRP is still the major cause for downgrading of seed potatoes in the Netherlands (Hiddink, 2022). The application of bacterial biocontrol agents could be a promising approach to control disease incidence and spread. Still, there is no widespread use of bacterial antagonists in potato cultivation.

The potential of biocontrol agents has been widely investigated in recent years, due to increased regulation of chemical pest protection and spread of diseases with no available protection. Strains from several bacterial species have shown effectiveness in pathogen control, such as strains from the genera *Bacillus*, fluorescent *Pseudomonas*, *Serratia*, *Burkholderia* and *Paenibacillus* species (Compant, et al., 2005). Mechanisms of antagonism include direct mechanisms such as competitive colonization of plant organs and the production of defensive compounds, including siderophores, lytic enzymes and antibiotic substances (Compant, et al., 2005, Raaijmakers, et al., 2002). Furthermore, some bacterial species can act against pathogens via indirect mechanisms, one prominent being the induction of resistance in the plant by priming systemic defenses (Pieterse, et al., 2014). Especially, species of the genus *Pseudomonas* have frequently been found to be effective biocontrol agents, as they are able to quickly colonize plants, are known to produce a wide range of siderophores and antibiotic compounds and have even been demonstrated to be involved in induced systemic resistance (ISR) in the plants (Weller, 2007).

A number of studies reported investigating biocontrol agents against SRP in potato tubers. Sturz and Matheson (1996) found *Curtobacterium luteum* to inhibit *P. atrosepticum* on solid cultivation medium, Krzyzanowska, et al. (2012) identified *Pseudomonas* species as potential antagonists in a tuber maceration assay, and Zhang, et al. (2020) assessed the ability of an *Acinetobacter* strain to reduce tissue maceration caused by *P. carotovorum*. In addition, it was found that *Serratia plymuthica* could reduce blackleg symptoms in the field caused by *Dickeya solani*, even if inoculated before storage (Czajkowski, et al., 2012, Hadizadeh, et al., 2019). It could also be shown that mixtures of different disease suppressive strains performed better than single strains in maceration assays and in greenhouse experiments (Krzyzanowska, et al., 2019, Raoul des Essarts, et al., 2016). However, it has frequently been found that results from *in-vitro* experiments on biocontrol are difficult to replicate in the field. This is due to high variability in abiotic conditions in the field, the presence of a high amount of other microorganisms and consequentially limited survival and activity of antagonists (French, et al., 2021). In addition, predicted antagonism on the basis of genomic features or *in vitro* assays might not be predictive of antagonism in the field. Therefore, field trials under realistic conditions should be considered at an early stage in the search for biocontrol agents. For potato, field trials with antagonistic microorganisms are scarce and to our knowledge no biocontrol organisms have been identified yet that provide consistent reduction of blackleg disease in the field.

The limited ability of *in-vitro* experiments to predict effectiveness in the field is mostly ascribed to lack of survival and plant colonization of the used biocontrol agents (Bashan, et al., 2014). Limited survival can be caused by adverse abiotic conditions or interactions with the indigenous microbial community (Qiu, et al., 2019). Moreover, introduced biocontrol agents need to be compatible with the specific plant species or even variety in order to establish in or around the plant. In their review Qiu, et al. (2019) suggest that the use of indigenous strains that are isolated from the specific environment that they are to be applied to, can overcome the issues described above. In addition, it has been found repeatedly that mixes or “synthetic communities” of biocontrol strains as opposed to single strains might be more effective (Krzyzanowska, et al., 2019). For one, it can be expected that differences in environmental preferences of different strains will ensure the survival of at least a subset of strains under changing conditions. Second, different bacterial taxa often possess different mechanisms of biocontrol and thereby can act in a complementary way to prevent or reduce disease. Thus, the selection of biocontrol agents that are naturally prevalent in potato tuber tissue and that can be combined in synthetic communities might be an effective strategy to generate effective antagonist mixtures.

In this study, we tested a range of potential biocontrol strains and combinations of them for their ability to reduce disease incidence of black leg in the field in four consecutive years. For this, we treated tubers infected with *D. solani* or *P. brasiliense* with potential antagonistic bacterial taxa isolated from potato plants. To ensure that the applied strains were still present on the day of planting, we tested survival of the antagonistic strains on potato tubers in storage. In addition, to uncover the potential mechanisms of antagonistic activity, for a selection of strains, we analysed whole genome sequences for the presence of genes related to antimicrobial production.

## Material and Methods

### Bacterial isolates

The 98 bacterial strains used in this study have been isolated previously in the project “Deltaplan Erwinia”. Isolates were obtained from mother tubers, seed tubers, stems and from soil surrounding tubers (Table S1). Selection of these isolates from in total 1344 isolates was based on the following criteria: *in-vitro* activity against *D. solani* (Ds), *P. brasiliense* (Pb) or *P. parmentieri* (Pp), presence in tubers and stems in high concentrations, the ability to macerate potato tissue (for quick degradation of the mother tuber, which reduces the risk of soft rot bacteria spreading to the rest of the plant), risk group 1 (non-pathogenic to humans or animals), and choice of a taxonomically diverse panel of strains. Of the selected strains, 72 were positive for *in vitro* antagonism against one or more of the three tested species (Ds, Pp and Pb).

In 2018 and 2019, 66 isolates were used. In 2020 and 2021, the isolates and mixes that had performed best in the previous years were used as well as isolates that had not been used previously, based on the same criteria.

### Experimental setup

For an overview of the experimental setup, see Figure 1.

**Fig. 1.**
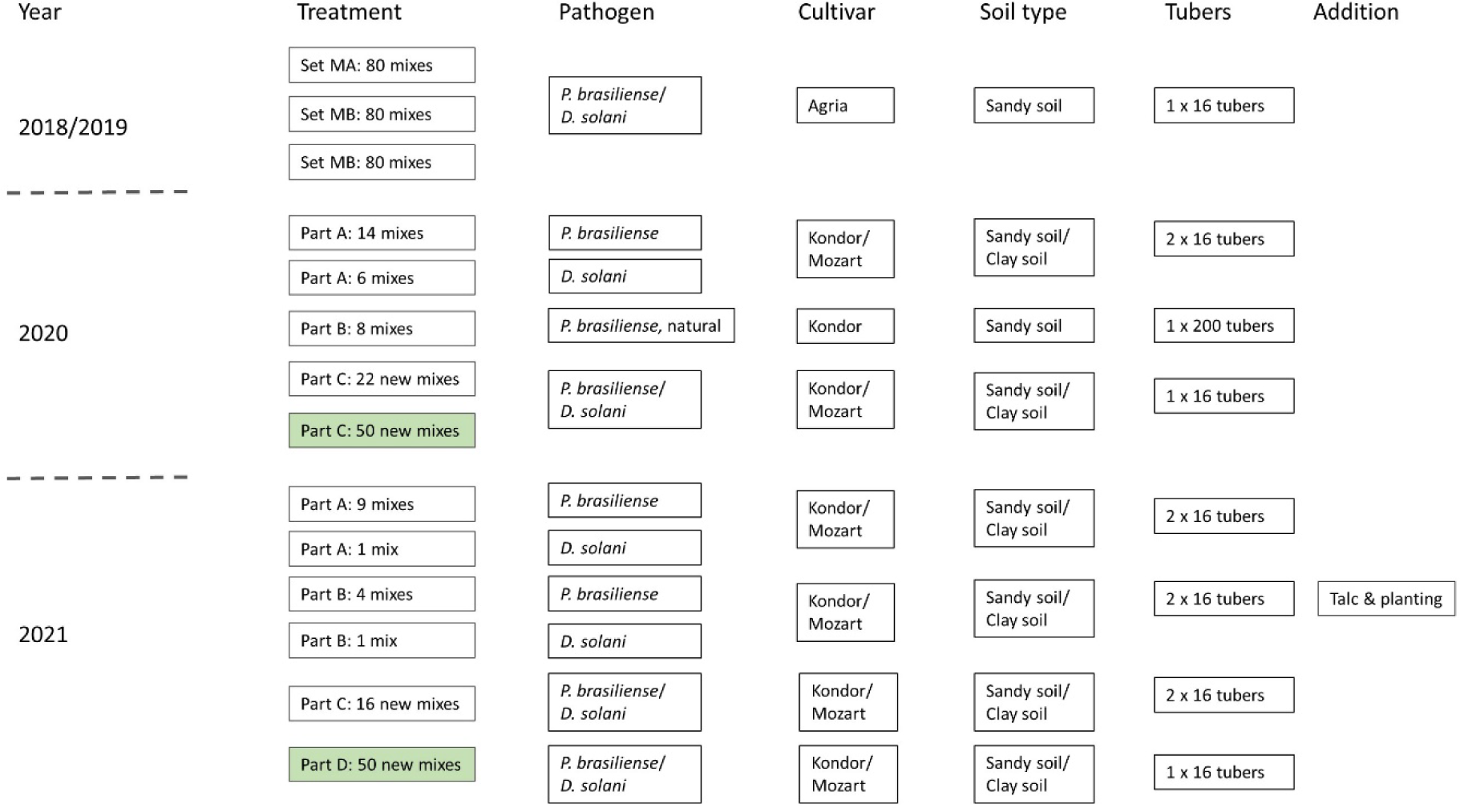
Experimental setup in the four years 2018, 2019, 2020 and 2021 with number of antagonist mixes used as treatments (treatments in light green consistent of new isolates not used in a previous year), inoculated pathogen, potato cultivar, soil type the tubers were planted in and number of tubers. Additional inoculation methods are indicated

#### Year 2018 and 2019

In 2018 and 2019, 66 strains (Table S1) were assigned to three different sets of 22 isolates each, called MA, MB and MC. Per set, these 22 strains were tested either alone or in mixes of three, yielding 80 different treatments per set and thus, 240 treatments in total excluding the water control (Table S2). All 240 treatments were tested against *D. solani* and *P. brasiliense*. Each treatment was used to inoculate 16 tubers of cultivar Agria per plot. The plots were planted in a field with sandy soil (Marknesse, The Netherlands) in a randomized order.

#### Year 2020

In 2020, 52 strains (Table S1) were used.

Part A consisted of in total 20 treatments which had been used in the two previous years (Table S3). Twelve of these mixes have been shown to be effective against *P. brasiliense*, whereas two have been shown to be ineffective in 2018 and 2019. These 14 mixes were inoculated on *P. brasiliense* infected tubers. Four of the mixes in part A have been shown to be effective against *D. solani*, whereas two were not effective. These six mixes were inoculated on *D. solani* infected tubers. Two cultivars, Kondor and Mozart were used. The treated tubers were planted in plots of 16 tubers, on two locations, sandy soil (Marknesse, The Netherlands) and clay soil (Munnikezijl, The Netherlands). Part A was planted with two replicate plots, resulting in 128 tubers per treatment.

In part B, eight of the mixes that were effective against *P. brasiliense* in 2018 and 2019, were additionally inoculated on naturally *P. brasiliense* infected tubers of the cultivar Kondor with 200 tubers per treatment (part B) (Table S3) and planted in sandy soil (Marknesse, The Netherlands).

In part C, 10 treatments contained strains used previously in 2018 and 2019, but combined in different mixes, 12 treatments contained strains used previously, but combined in mixes of two strains and 50 mixes contained strains not used previously (Table S3). Tubers were inoculated either with *D. solani* or *P. brasiliense*. Also, for this part the cultivars Mozart and Kondor were used in plots of 16 on the two locations. Part C was planted with one replicate, resulting in 64 tubers per treatment.

#### Year 2021

In 2021, 55 strains were used (Table S1).

In part A, the ten most effective treatments from the previous years were used, one for *D. solani* and nine for *P. brasiliense* (Table S4). These were again inoculated onto the cultivars Kondor and Mozart and planted in sandy and clay soil, in plots of 16 with two replicates, resulting in 128 tubers per treatment.

In part B, four of those treatments were chosen (one for *D. solani* and three for *P. brasiliense*) and either inoculated with a talc formulation or by direct inoculation in the planting hole. The same number of tubers as in part A was used.

In part C, the 25 strains that occur in the ten treatments used above in part A, were combined in 16 new mixes of five strains each and inoculated in a similar manner on tubers infected with both pathogens (part C), resulting in 128 tubers per treatment.

In part D, 20 new isolates were combined in 50 treatments and inoculated on Mozart and Kondor tubers infected with both pathogens. These were planted with only one replicate in both sandy and clay soil, resulting 64 tubers per treatment.

In all years, positive controls were included that were inoculated only with one of the two pathogens and water controls, inoculated only with water, with four replicates per pathogen, cultivar and location resulting in 64 tubers per control in 2018 and 2019 and 256 tubers per control in 2020 and 2021.

### Tuber inoculation

All tubers, with exception of the water control, were vacuum inoculated with either *P. brasiliense* or *D. solani* at a concentration of 10^6^ cells/ml at HZPC (Metslawier, The Netherlands). Every year, 15 tubers of each cultivar inoculated with each of the pathogens were used for assessing the inoculation efficiency. For each tuber, the peel was crushed, extraction buffer (Ringer’s buffer with 0.1% DIECA) was added at twice the peel weight and the extract was plated on TSA for colony counting.

At Wageningen University and Research (Wageningen, The Netherlands), two days prior to inoculation with the antagonists, the antagonist strains were plated on TSA medium. After two days of incubation at 25°C the isolates were scraped from the plates and suspended in 4 ml Ringer’s buffer per plate, resulting in a dense solution of approximately 7×10^9^ cells/ml. To prepare single strain inoculums and mixes, 2 ml of each strain in a mix was used and diluted with tap water to 50 ml, resulting in an inoculum with a concentration of 10^8^ cells/ml (note that mixes with three strains contain three times as many cells as treatments with only one strain). The inoculums were transferred to plant spray bottles and sprayed on the tubers under sterile conditions, using 50 ml per 32 tubers. The tubers were then air-dried, packed in paper bags and stored at 4°C for approximately two weeks before planting.

In 2021, four treatments were used as a talc formulation. These formulations were prepared according to Dhar Purkayastha, et al. (2018). In addition, the same four treatments were inoculated on tubers in the planting hole by pouring 50 ml of the inoculum on each tuber after planting.

The tubers were planted at the beginning of May and the plants were checked biweekly for symptoms of blackleg until the beginning of July.

### WGS sequencing

In 2020, DNA from 78 antagonist isolates was extracted using the sbeadex Agowa kit according to manufacturer’s instructions. Nextera XT library preparation and Illumina HiSeq of 350 bp paired end reads was done at BaseClear (Leiden, The Netherlands).

Adapter trimming and de novo assembly of the reads was done in the CLC Genomics Workbench 20. Six strains that were assembled into short contigs with CLC were again assembled with SPAdes 3.14.1.. An average nucleotide identity (ANI) analysis was carried out using OrthoANI. Subsequent steps were only performed for the 25 strains that were used in the 10 treatments in part A of the field experiment in 2021. All scaffold genomes were analysed with BUSCO v.5. In addition, a taxonomic analysis was done using GTDB-Tk (Chaumeil, et al., 2019).

In order to identify genes that are potentially involved in antagonistic activity, all scaffold genomes were submitted to Antismash v. 5 (Blin, et al., 2019). The first hits of the KnownClusterBlast results were used if similarity was ≥50%.

### Primer design and PCR reaction conditions

Primers for PCR assays were designed for the 25 isolates that were used in 2021 from the ten most effective treatments of the previous years. These assays were developed to be specific for each isolate in a background of the 24 other isolates. The ANI analysis identified the isolates MT096 and MT099 as identical and therefore they share a primer pair. Design was done in CLC Genomics Workbench 20. For all primer sequences see table S5.

Reactions consisted of 6 µl GoTaq buffer (5×), 0.6 µl Forward primer and Reverse primer each (10 µM), 1.2 µL dNTPs (5 mM), 0.15 µl GoTaq G2 and 19.45 µl DNAse-free water with the addition of 1 µl sample DNA or briefly heated single colony. The assay was run under the following conditions: 2 min at 94°C, followed by 35 cycles of 15 sec at 94°C, 20 sec at 55°C, 45 sec at 72°C. The final step was 5 min at 72°C. Only the assay for isolate C5 required an annealing temperature of 60°C. All assays were tested for specificity.

### Survival in storage

In order to assess if the antagonist strains survive on the tubers in the period of two weeks from inoculation to planting, survival of nine single strains and the ten most effective mixes (Table S4, Part A) was tested. For single strains, tubers of the cultivar Axion were used and for mixes the cultivar Frieslander. Tubers were surface sterilized according to Czajkowski, et al. (2012). Subsequently, inoculation was done similar to inoculation with antagonists as described above. Tubers were sprayed with water as a negative control and dried and stored at 4°C in sterilized boxes. Samples were taken after 2, 4, 7, 10, 14 and 15 days. At each day with the exception of day 15, three tubers with identical treatments were peeled and the peels were crushed with 2 ml Ringer’s buffer in BioReba bags in a hammering machine. Serial dilutions of 1:10 to 1:1000000 were prepared in Ringer’s. Of each dilution, 100 µl were mixed with 300 µl liquid LB agar at a temperature of 48°C, containing 0.3 µl cycloheximide. The agar was allowed to solidify, and the plates were incubated at 25°C until colony formation. Colonies from each dilution were picked with sterile toothpicks, dissolved in water, and heated for 5 min at 90°C. These samples were stored at -20°C until PCR analysis. PCR analysis for the individual strains was carried out to determine at which day and which dilution the target strains could still be detected. At day 15, one tuber from each treatment was exposed to UV-light in a plant growth chamber before sampling. This was done to simulate exposure to sunlight in the field before planting.

### Survival in the field

In order to assess if the used antagonists have survived and multiplied in the plant throughout the growing season, at the end of the experiment in July in 2021, stems were harvested from three treatments (4, 19, 24, see Table S4) from the cultivar Mozart in the clay soil. Stems from the water control and the positive controls were sampled as well. For each treatment, in total six plants were sampled, including healthy as well as diseased plants if available. Only healthy stems were sampled by cutting of the first 10 cm from the ground and cleaning them of soil. They were cooled and transported immediately to WUR for further processing.

Stems were surface sterilized by immersion in 0.05% hypochlorite solution for 1 min, immersion in 70% ethanol for 1 min and 3 times washing in tap water. The stems were cut into pieces and 5-15 g of stem tissue with an equal amount of Ringer’s buffer was placed in a Bioreba-bag and crushed with a hammering machine. One ml of homogenized extract was transferred to an Eppendorf tube, centrifuged and the supernatant was removed. The pellets were stored at -20°C until DNA extraction.

DNA extraction was done with the sbeadex Agowa kit using the mini-protocol on a Kingfisher robot. PCRs were done with the primers designed previously. Each treatment was tested with all the primers designed for the strains that have been inoculated into the respective treatment. Controls were tested with all primers used in this experiment to detect potential non-specific amplification. For each strain, cells from the respective isolate were used as a positive control.

### Statistical analysis

#### 2018 and 2019

To assess if pathogen or treatment (antagonist, positive control, negative control) had an effect on disease incidence a generalized linear model was used with binomial distribution and a logit link. For pairwise comparison of the treatments the emmeans function from the package emmeans was used (Lenth, et al., 2018). Similarly, it was tested if there was a difference in disease incidence between treatments with only one antagonist strain and treatments consisting of a mix of three strains.

#### 2020

To determine significant differences between treatments regardless of location and cultivar a glmmPQL from the package MASS (Venables and Ripley, 2013) was used, with location and cultivar as random variables. In part C, for *P. brasiliense* and *D. solani* separately, treatments were compared with a general linear model. For pairwise comparison of the treatments the emmeans function was used. The effects of location and cultivar on disease incidence was tested using a generlized linear model with binomial distribution and a logit link. If interactions occurred, those were tested pairwise using the emmeans function.

#### 2021

To determine significant differences between treatments regardless of field pathogen and cultivar, a glmmPQL from the package MASS (Venables and Ripley, 2013) was used, with location, pathogen and cultivar as random variables. For each part, except D, general linear models (glm) were performed with treatment, location, and cultivar as explanatory variables, separately for tubers treated with *P. brasiliense* and *D. solani*. For pairwise comparison of the treatments the emmeans function was used.

A spearman correlation test was performed between disease incidence in the 10 mixes of 25 antagonists used in 2020 and 2021 and the number of different compounds per mix as identified by AntiSmash.

All statistical analysis was carried out with RStudio (2016).

## Results

### Year 2018 and 2019

In 2018, emergence from planted tubers was generally low with on average only 32% of all planted tubers emerging, while in 2019 emergence was on average 97%. Therefore, the experiment in 2018 can be viewed as failed and was not taken further in the analysis of results. In 2019, disease incidence was 40% higher in tubers inoculated with *P. brasiliense* than with *D. solani* (z-ratio=-10.13, p<0.01). On average, antagonist treatment led to a reduction of disease incidence compared with the positive control for both *P. brasiliense* and for *D. solani* (Table 1). The water control showed no disease. Several treatments were effective in reducing disease incidence, but effects of treatments differed between *D. solani* and *P. brasiliense* (Figure 2). For tubers inoculated with *P. brasiliense*, disease incidence was on average 25% higher in treatments with single strains compared to mixes (z-ratio=-8.12, p<0.01; data not shown).

**Table 1:** Statistical results of a comparison of disease incidence between the antagonist treatment and the positive and negative control in the years 2018 and 2019.

| Year | Pathogen | Comparison | Disease incidence antagonist | Disease incidence control | z-ratio | p |
| --- | --- | --- | --- | --- | --- | --- |
| 2019 | *D. solani* | Antagonist-positive control | 46.89±12.22 | 59.38±11.69 | -2.55 | 0.03 |
| 2019 | *D. solani* | Antagonist-water control | 46.89±12.22 | 0 | 0.02 | 0.99 |
| 2019 | *P. brasiliense* | Antagonist-positive control | 61.00±29.78 | 85.89±10.27 | -4.46 | <0.01 |
| 2019 | *P. brasiliense* | Antagonist-water control | 61.00±29.78 | 0 | 0.05 | 0.99 |

**Fig. 2.**
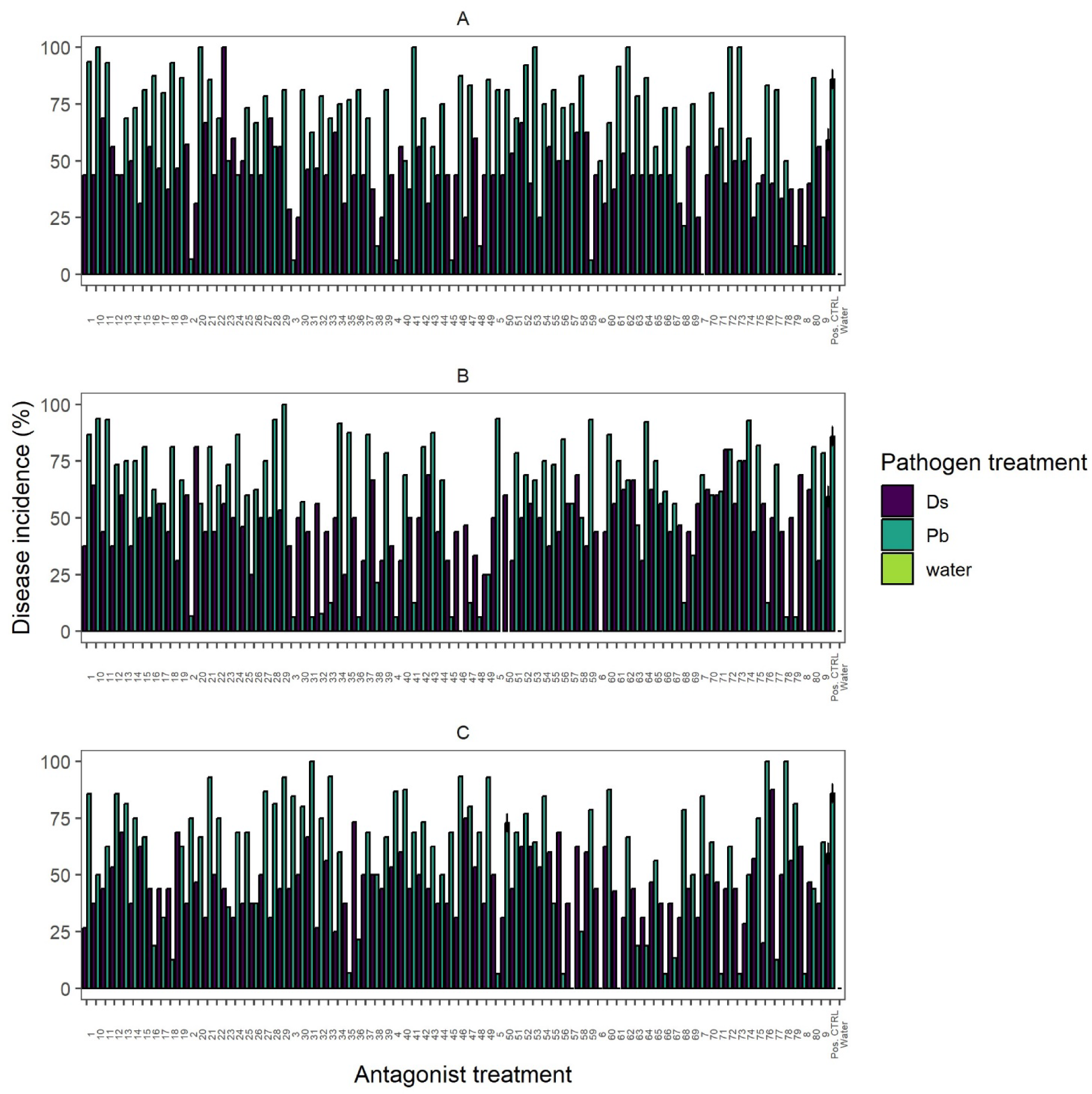
Disease incidence with the different antagonist treatments in the three sets MA, MB and MC and in the positive and water control in 2019 for a) *D. solani* and b) *P. brailiense*; positive and water controls are identical for all sets; error bars represent the standard error

### Year 2020

In 2020 part A, 20 treatments that had been used previously in 2019 were applied again. Fourteen of those were previously effective against *P. brasiliense* and four against *D. solani* (Table S3). Two each had previously shown no disease reduction against the respective pathogen in 2019. In addition, all treatments were tested on two locations (sandy soil and clay soil) and two potato cultivars (Mozart and Kondor), with two replicates each.

Unlike in 2019, no treatment in 2020 part A showed a significantly different disease incidence from the positive control (data not shown). Also, disease incidences differed from the results from the previous year (Fig. 3). However, location and cultivar had effects on disease incidence. For tubers inoculated with either pathogen, disease incidence was significantly higher in cultivar Kondor than in cultivar Mozart and higher in sandy soil than in clay soil (Fig. S1, for statistical results see Table S5). The effects of treatment on disease incidence varied between locations and cultivars (tubers in inoculated with *P. brasiliense*: χ^2^=37.35, p<0.01; tubers inoculated with *D. solani*: χ^2^=20.37, p<0.01), but at no location and in no cultivar did treatment with antagonists reduce disease incidence compared to the positive control (data not shown).

**Fig. 3.**
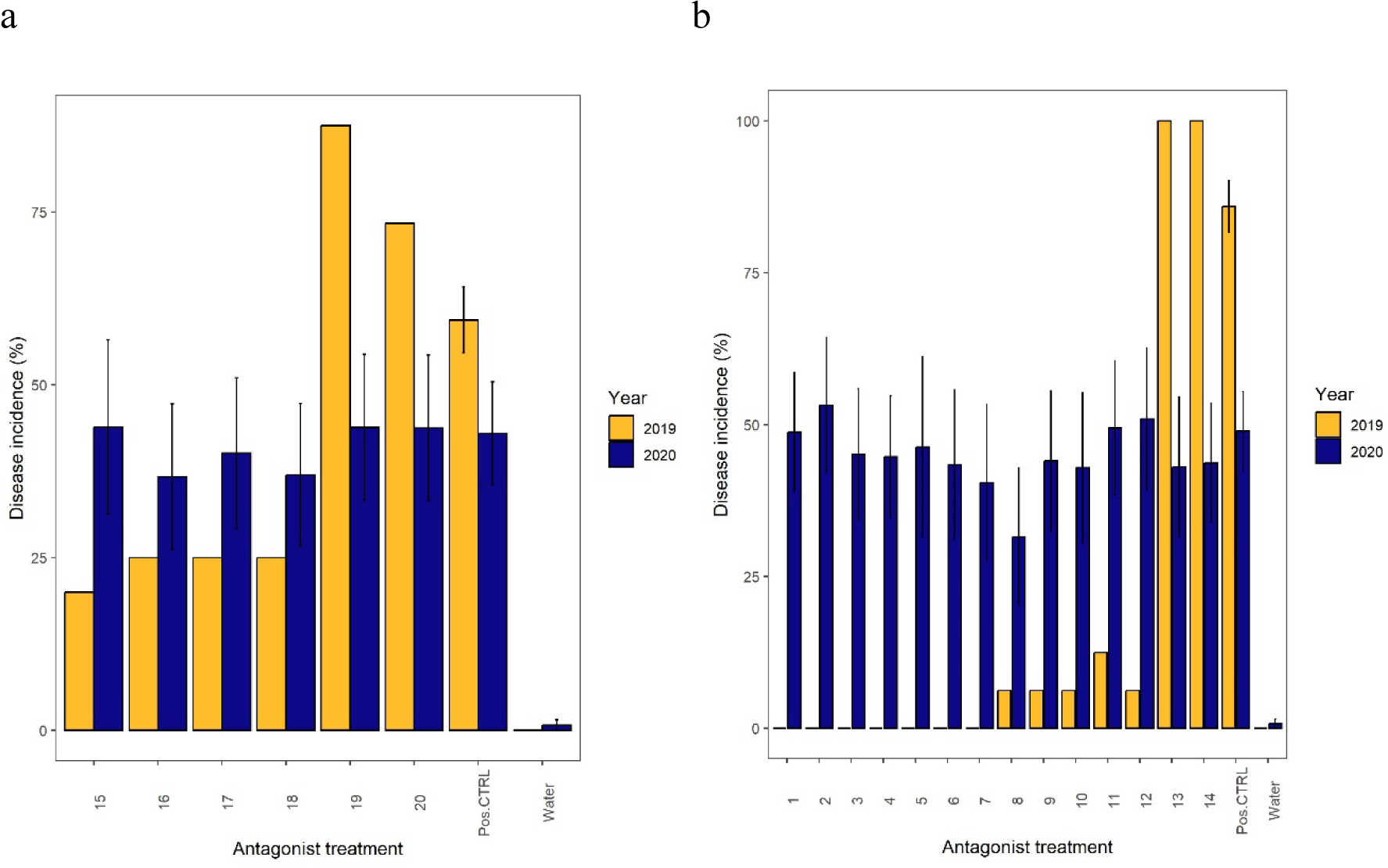
Disease incidence in the different antagonist treatments used in 2020 part A and 2019; in 2020 averaged over location, cultivar and replicate for a) *D. solani* and b) P*. brasiliense*; error bars represent the standard error

In part C, new mixes of antagonists were tested on tubers infected with either *D. solani* or *P. brasiliense* (Table S3).

Similar to part A, disease incidence was higher in cultivar Kondor and in sandy soil (for statistical results see Table S5) and treatment effect varied with both location and cultivar (tubers in inoculated with *P. brasiliense*: χ^2^=228.03, p<0.01; tubers in inoculated with *D.solani*: χ^2^=235.83, p<0.01) (data not shown). No treatment led to a reduction in disease incidence compared to the positive control for either *P. brasiliense* or *D. solani* when averaged over cultivar and location (Fig 4). However, if only sandy soil location was analyzed, disease incidence with *P. brasiliense* was significantly lower in the treatments 37, 68, 127 and 130 compared to the positive control (Table S6). In addition, in cultivar Kondor inoculated with *P. brasiliense* disease incidence was lower in the treatments, 34, 35, 68, 69, 129, 130 and 131 compared to the positive control (Table S6). For tubers inoculated with *D. solani*, no treatment decreased disease incidence in either location or cultivar.

**Fig. 4.**
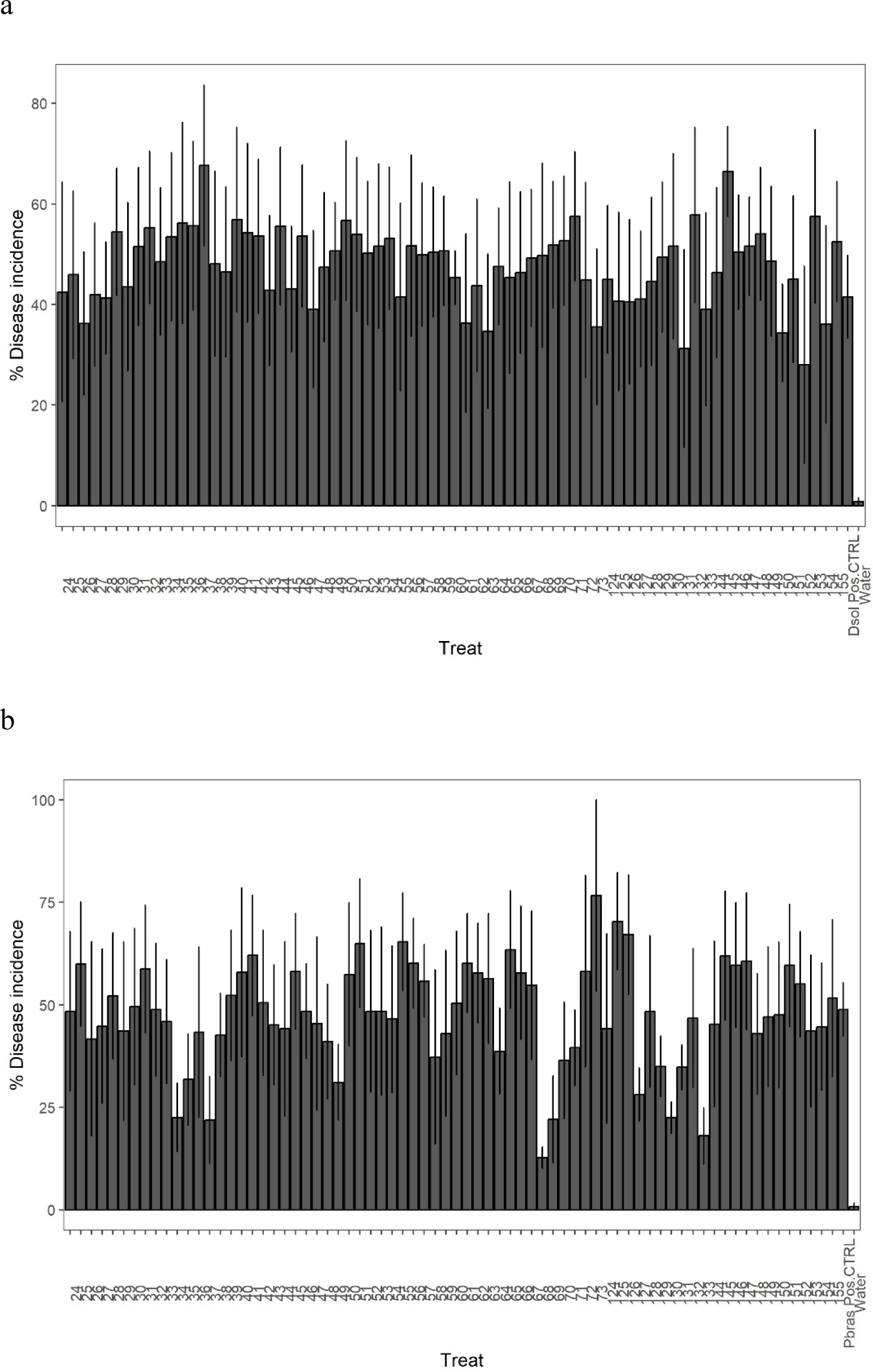
Disease incidence in plants with the different treatments averaged over location, cultivar and replicate, for plant inoculated with a) *D. solani* and b) *P. brasiliense*; error bars represent the standard error

In part B, tubers that were naturally infected with *P. brasiliense*, disease incidence was very low, with an average of 5%. Thus, differences between treatments were small (Fig. S2).

### Year 2021

In 2021 part A, ten treatments were used that had been successful in reducing disease incidence in 2020 (Table S4). Nine of these treatments had been moderately effective against *P. brasiliense* and one against *D. solani*. Those were again inoculated on the two races Mozart and Kondor and planted in sandy and clay soil.

Disease incidence was higher on average in cultivar Kondor compared to cultivar Mozart and higher in sandy soil, but only for cultivar Mozart (Table S7).

There was an effect of treatment on disease incidence on average (χ2=409.26, p<0.01), but there were no pairwise differences between any of the single treatments and the positive controls (data not shown). In addition, there were interactions between cultivar and treatment (χ2=27.65, p=0.09) and location and treatment (χ2=45.27, p<0.01) (data not shown). However, for none of the locations and cultivars a significant reduction of one of the treatments compared to the positive controls was detected. While the most effective treatments from 2020 were selected, disease incidence was significantly higher in 2021 with 45% average disease incidence compared to 2022 with 35% average disease incidence (χ2=40.46, p<0.01). This was the case for almost all treatments except treatment 1 for *D. solani* and treatment 3, 4 and 10 for *P. brasiliense* where the difference between 2020 and 2021 was not significant (Fig. 5).

**Fig. 5.**
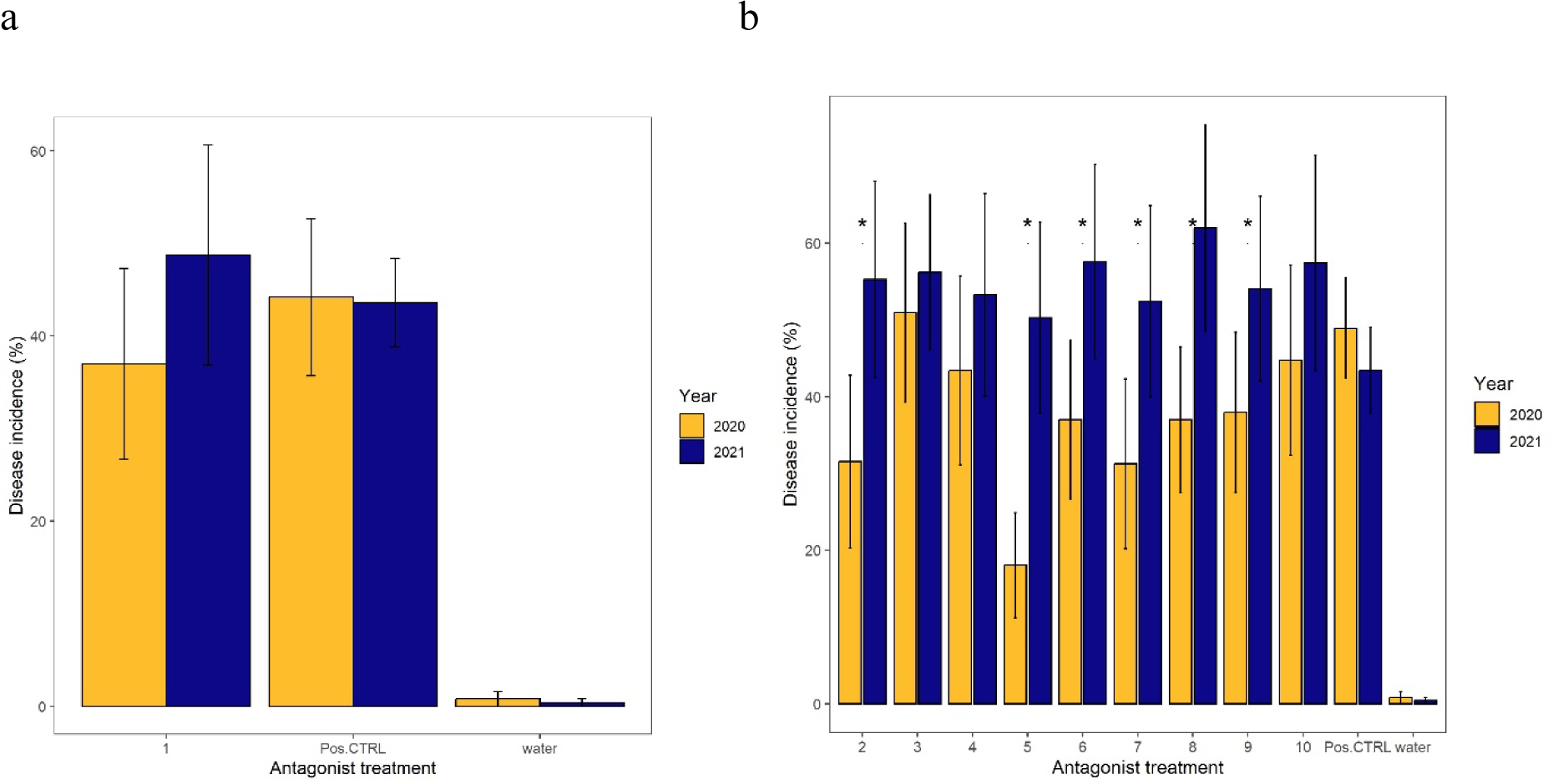
Disease incidence in 2020 and 2021 for plants treated with antagonists and in the positive and water control for a) *D.solani* and b) *P. brasiliense*; error bars represent the standar error

In part B, four treatments also contained in part A were tested as a talc formulation (B1) and when applied directly into the planting hole (B2). In both cases, we observed again a higher disease incidence in Kondor than in Mozart and in sandy soil compared to clay soil, but only for cultivar Mozart (Table S7). No differences could be observed between the treatments and the positive controls with respect to disease incidence (data not shown).

In part C, there was also an overall higher disease incidence in cultivar Kondor compared to Mozart (χ^2^=811.70, p<0.01) and sandy soil compared to clay soil (χ^2^=74.69, p<0.01), except for disease incidence in potatoes of cultivar Kondor inoculated with *P. brasiliense*, which is higher in clay soil than in sandy soil (Table S7). Also in this part, no treatment significantly decreased disease incidence compared to the positive control, neither for tubers inoculated with *P. brasiliense* nor with *D. solani* (Fig. 6). There were interactions between cultivar and treatment (χ^2^=68.60, p<0.01) and location and treatment (χ^2^=33.38, p<0.01), but at no location and no cultivar any treatment decreased disease incidence.

**Fig. 6.**
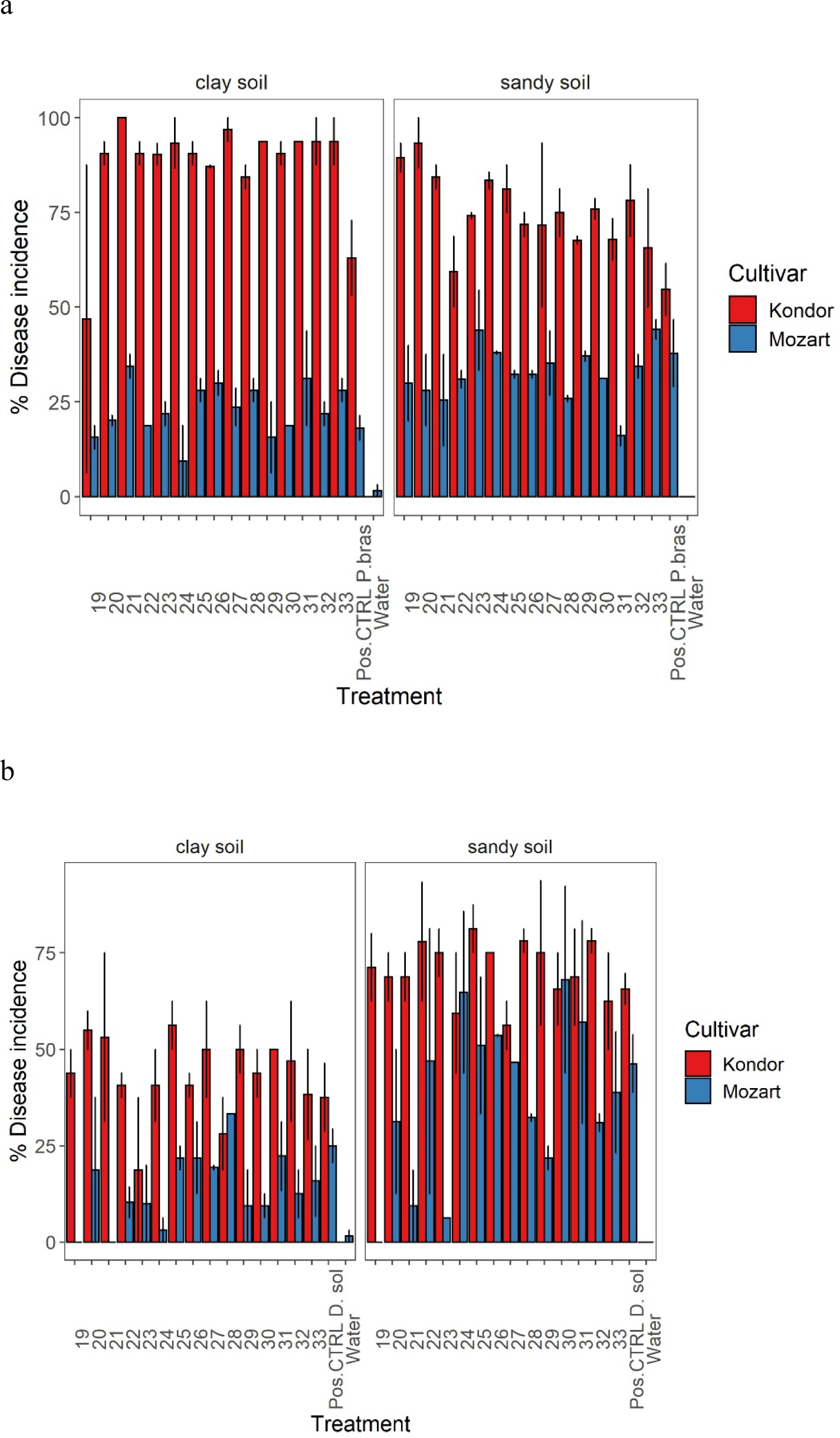
Disease incidence in the treatments in part C in cultivar Kondor and Mozart in sandy and clay soil for tubers inoculated with a) *D. solani* and b) *P. brasiliense*; error bars represent the standar error

In part D, again disease incidence was higher in cultivar Kondor compared to cultivar Mozart (χ^2^=703.37, p<0.01) and higher in sandy soil compared to clay soil for cultivar Mozart (Table S7). None of the treatments decreased disease incidence compared to the positive control (data not shown).

### WGS

All 78 isolates that were sequenced were successfully *de novo* assembled. An ANI analysis showed that eleven strains were highly identical (Fig. S3). For the 25 selected strains (Table S1), the taxonomic identity was determined with GTDB-Tk. Furthermore, they were analysed with BUSCO to assess completeness of the genome and to detect potential contaminations (Table S8). The genomes showed a high completeness. In strain A11, a contamination with *Pantoea vagans* could be detected.

Of all 78 analysed strains, in 65 one or more genes were detected coding for a potential antimicrobial compound (data not shown). Here we focused on the 25 strains in the ten treatments used in 2021 part A. Of these 25 strains, 22 possessed genes with potential antagonistic functions (Table 2). Most of these compounds are non-ribosomal peptides (NRPs) produced by *Pseudomonas* species. These compounds include bananamide, viscosin, lokisin, fragin and tolaasin. Five strains were potentially able to produce siderophores. The strains, C5 (*Stenotrophomonas rhizophila*), N2 (*Paenibacillus polymyxa*) and N3 (*Bacillus pumilus*) were notable as they harboured genes involved in the production of 4-6 different compounds. Since many compounds were unique to one isolate, whereas others were produced by a great number of isolates, it was not possible to test if the presence of specific compounds was correlated with disease incdence in plants inoculated with these strains. However, there was no correlation between the number of different compounds per treatment and the disease incidence in 2020 and 2021 (rho=0.02, p=0.84). Notably, some strains for which no genes for antimicrobial compound production were found, still had previously shown *in vitro* antagonism against SRP in 2019 when single strains were tested.

**Table 2:** Presence of genes that code for potentially antimicrobial compounds; only genes with a similarity of >50% or higher were taken into consideration

| Strain ID | ID | Species | Genes | Functions |
| --- | --- | --- | --- | --- |
| A5 | H007 | *Pseudomonas fluorescens* | bananamide (NRPS),<br>viscosin (NRPS) | banamide=antifungal,<br>viscosin= antibacterial |
| A1 | H017 | *Pseudomonas koreensis* | lokisin (NRPS) | lokisin=antifungal |
| C5 | H027 | *Pseudomonas* sp. | tolaasin (NRPS),<br>viscosin (NRPS),<br>pyochelin (NRPS),<br>pyocanine (phenazine) | tolaasin=antibacterial,<br>viscosin=antibacterial,<br>pyochelin=siderophore,<br>pyocyanine=antibacterial |
| A6 | H045 | *Pseudomonas putida* | lokisin (NRPS),<br>fragin (NRPS-like) | lokisin=antifungal<br>fragin=antibacterial |
| A3 | H099 | *Pseudomonas koreensis* | lokisin (NRPS),<br>rhizomide (NRPS) | lokisin=antifungal,<br>rhizomide=antibacterial |
| A11 | H100 | *Pseudomonas marginalis*+<br>*Pantoea vagans* | tolaasin (NRPS) | tolaasin=antibacterial |
| B1 | H110 | *Pseudomonas* sp. | bananamide (NRPS) | banamide=antifungal |
| C1 | MT086 | *Stenotrophomonas* sp. |  |  |
| N8 | MT096 | *Pseudomonas* sp. | lokisin (NRPS) | lokisin=antifungal |
| C2 | MT099 | *Pseudomonas* sp. | lokisin (NRPS)<br>fragin (NRPS-like) | lokisin=antifungal<br>fragin=antibacterial |
| C18 | MT251 | *Curtobacterium*<br>sp. | carotenoid (terpene),<br>desferrioxmin B<br>(siderophore) | carotenoid: protection<br>from oxidative stress,<br>desferrioxmin B=<br>siderophore |
| C20 | MT293 | *Pseudomonas* sp. | fragin (NRPS-like) | fragin=antibacterial |
| N1 | MT330<br>a | *Paenibacillus* sp. | paeninodin<br>(lassopeptide) |  |
| A21 | MT453<br>b | *Serratia*<br>*fonticola* | aryl polyene (aryl<br>polyenes), aerobactin<br>(siderophore) | aryl polyene= protection<br>from oxidative stress,<br>aerobactin=siderophore |
| N5 | MT518<br>b | *Pseudomonas* sp. | lokisin (NRPS)<br>fragin (NRPS-like) | lokisin=antifungal<br>fragin=antibacterial |
| N4 | MT546<br>a | *Pseudomonas* sp. | lokisin (NRPS), 3-thiaglutamate (PRE-containing, RiPP-like) | lokisin=antifungal |
| N2 | MT558<br>a | *Paenibacillus*<br>*polymyxa* | paenilan<br>(lanthipeptide),<br>anabaenopeptin<br>(NRPS),<br>paenilopoheptin<br>(NRPS), fusaricidin<br>(NRPS), tridecaptin<br>(NRPS), polymyxin<br>(NRPS),<br>paenilipoheptin (NRPS) | paenilan= antibacterial,<br>fusaricidin=antobacterial,<br>tridecaptin =antibacterial,<br>polymixin=antibacterial |
| N12 | MT580<br>a | *Pseudomonas* sp. | lokisin (NRPS) | lokisin=antifungal |
| N3 | S022 | *Bacillus altitudinis* | caroteinoid (terpene),<br>lichenysin (NRPS),<br>bacillibactin (NRPS),<br>fegycin (betalactone), | lichenysin= antobacterial,<br>bacillibactin=siderophore,<br>fegycin=antibacterial |
| B7 | S035 | *Pseudomonas fluorescens* | fragin (NRPS-like)<br>pseudomonine (NRPS) | fragin=antibacterial<br>pseudomonine=siderophore |
| C16 | S039 | *Pseudomonas fluorescens* | pseudomonine (NRPS)<br>fragin (NRPS-like) | pseudomonine=siderophore<br>fragin=antibacterial |
| C3 | T032 | *Pseudomonas* sp. | fragin (NRPS-like) | fragin=antibacterial |
| N11 | T083 | *Staphylococcus vitulinus* |  |  |
| C15 | T091 | *Sphingobium yanoikuyae* | carotenoid (terpene),<br>ectoine (ectoine) | carotenoid= protection<br>from oxidative stress,<br>ectoine= protects cell<br>membrane from surfactant |
| N10 | T150 | *Staphylococcus* sp. | staphylobactin<br>(siderophore),<br>staphyloferrinA<br>(siderophore) | staphylobactin=siderophore,<br>staphyloferrinA=siderophore |

### Survival in storage

Of the eight isolates tested for survival on tubers in storage as single strains, seven isolates could be detected after 14 days and 15 days after exposure to light after diluting the extract from the peel 1:1000, which is equal to approximately 10^5^ cfu/ml (Fig. S4). Only isolate MT251 could not be detected in any dilution at any day. However, it must be noted that also the isolate kept on plate lost its ability to grow on TSA medium when transferred to a new plate.

In the mixes as well, 23 of 25 used isolates could be detected at the end of the storage time (Fig. S5). Again, MT251 could not be retreived as well as H045, which might also have been due to impaired growth on TSA medium. T150 could not be retrieved after exposure to light after 15 days.

### Survival in the field

Of all twelve isolates that had been inoculated in total in the samples taken from the field at the end of the growing season, one strain, C5 (*Pseudomonas fluorescens*) could be detected by PCR in one stem sample from a tuber previously treated with a mix of C5, C15 and C18 (treatment 4).

## Discussion

In this four-year study we used potential bacterial biocontrol strains obtained from potato tissue and rhizosphere soil to mitigate blackleg disease caused by *Dickeya solani* and *Pectobaterium brasiliense*. We found that several mixtures of biocontrol agents reduced blackleg disease incidence. However, this effect was not constant, but varied between years, locations, plant cultivars and replicates. In the following we will first discuss overall trends and then results from each of the four years.

In all years, there were considerable effects of pathogen, cultivar and location on disease incidence. A higher disease incidence was observed in plants of which the tubers had been inoculated with *P. brasiliense* compared to *D. solani*. This confirms the assumption that *P. brasiliense* is the more aggressive pathogen and might explain its rapid spread in many countries around the world (van der Wolf, et al., 2017). In addition, disease incidence was found to be constantly higher in cultivar Kondor than in cultivar Mozart in the years the two were compared. As no cultivar is known to be completely resistant to SRPs, not many published studies have focused on differences between resistance in cultivars. Nevertheless, current research has shown that these differences exist (Van Gijsegem, et al., 2021). Especially, the cultivars Agria, Kondor and Agata have been found to be susceptible to blackleg. For the first two cultivars, susceptibility was supported in our study where high disease incidences were constantly detected in the positive controls. We also found disease incidence to be higher on average in sandy soil compared to clay soil. It has been proposed that this might be due to higher temperatures in sandy soil compared to clay soil, favouring pathogen growth. However, the effect of soil type on blackleg disease has not yet been studied.

In 2018, poor emergence prevented us from conclusively determining treatment effects on disease incidence. As emergence was poor in the water control as well, we did not use emergence as a measure of symptom expression. In the following years, emergence was high regardless of treatment. In addition, in 2019 large differences between treatments could be observed. For *P. brasiliense*, a number of plots with treated plants showed no disease symptoms. It was observed that mixes of antagonists with three strains performed better on average than single strains. This confirms previous studies that showed the enhanced biocontrol efficiency of antagonist combinations (Krzyzanowska, et al., 2019). Combinations are assumed to be more effective due to the employment of several complementary antagonistic activities and a higher stability under changing conditions as different species have different niche optimums.

In the years 2020 and 2021, treatments that had been effective previously were repeated and complemented by the use of additional potential biocontrol strains. However, treatments that efficiently reduced disease incidence in the previous year did not show the same effect in the following year. Variation of biocontrol efficiency is frequently observed when using microorganisms as biocontrol agents. This variation is mostly attributed to variation in biotic and abiotic parameters between the years. Survival and spread of the biocontrol agents are likely to be dependent on factors such as the microbial community composition in the soil and tuber, temperature, moisture, and other soil properties (Amir and Alabouvette, 1993). While conditions can be favorable in one year and at one location, they can be adverse to the growth of the biocontrol strains at others. This is supported by the considerable variation between cultivars, locations and even replicates. The latter indicates that local conditions might play an important role. To increase antagonist survival in 2021, several treatments were repeated using a talc formulation or by supplying the bacterial inoculum to the tuber during planting. Talc formulations are frequently used to improve spread and adhesion of the biocontrol strains to the tuber (El-Hassan and Gowen, 2006). Application of biocontrol strain inoculums in the field reduces loss of living cells during storage and thereby possibly enhances bacterial concentrations. However, none of these treatments were more effective than spraying of the tubers. Also of the new mixes, used for the first time in 2020, only few showed some decrease of disease symptoms and this effect could not be reproduced in the following year.

In this study we attempted to achieve stable biocontrol activity by using bacterial strains that colonize the plants emerging from the tubers. Biocontrol strains were expected to enter through the lenticels becoming endophytic, where they are protected from environmental conditions and come in direct contact with the pathogen. Analysis of plant stems from treatments with a low disease incidence only detected one of the inoculated strains, H27, identified as *Pseudomonas fluorescens*, which has been isolated from potato stems. Strains of the genus *Pseudomonas* have been reported as endophytes in potato (Andreote, et al., 2009, Mercado-Blanco and Bakker, 2007). They have even been shown to alter the endophytic community and act as antagonists against diseases such as *Phytophora infestans* or *Fusarium oxysporum*. However, no constant efficiency against *D. solani* or *P brasiliense* could be found in the present study. The other strains tested for were not detected and might have failed to colonize the plant. Although most used strains were isolated from plant tissue, they might differ in colonization efficiencies are known. Some species might not be able to colonize the plant through the tuber, but enter through the rhizosphere (Diallo, et al., 2011). Moreover, Andreote, et al. (2010) found that colonization differs between potato cultivars and plant growth stage. Other methods to ensure efficient colonization might be the use of vacuum infiltration or addition of the antagonists after harvest and before storage.

As antagonist mixtures failed to elicit a consistent effect, we sought to confirm antagonistic potential by whole genome sequencing and analysing secondary metabolite production. Most obtained genomes were highly fragmented with a high number of small contigs, which might have prevented us from identifying all loci of interest. Nevertheless, most strains and especially members of the genus *Pseudomonas* harboured a number of genes that play a role in the production of antibacterial or antifungal compounds and genes for the production of siderophores. Siderophores are involved in iron sequestration and can therefore be effective in outcompeting pathogens in the uptake of iron (Leong, 1986). In addition, siderophores can also induce systemic resistance in plants (Bakker et al., 2003). The presence of these genes indicates the potential of the used isolates to elicit biocontrol. In spite of these identified gene loci, there was no correlation between the number of genes for secondary metabolite production in a mic of beneficial strains and reduction of disease incidece. It has to be taken into consideration that the presence of genes for the production of a certain metabolite does not automatically implicate the production of this compound *in-situ*.

We monitored antagonist survival in storage to ensure that living cells would remain at the time of planting. Most strains were still present after 14 days of storage and after a treatment with natural light. Detection limit was a maximal dilution of 1:100, which indicates concentrations of approximately 5×10^5^ cfu per tuber determined with semiquantitative plating. Thus, cell concentration decreased during storage. A previous study on the survival of bacterial antagonists in cold storage showed that a number of isolates did not survive well at low temperature (Hadizadeh, et al., 2019). Isolates that did survive, showed a similar decrease in concentration than was found in this study. Nevertheless, Hadizadeh, et al. (2019) found significant biocontrol of soft rot in storage, suggesting that survival was sufficient for biocontrol activity. In the present study, only two strains of all 25 strains, applied in ten mixes could not be recovered. This was likely due to slow or no growth on high nutrient media, such as LB and TSA, as also transfer of the same strains to fresh plates resulted in no or little growth. Attenuation of growth on common growth media is known as the viable but non-culturable (VBNC) state (Oliver, 2010). It is possible that all strains survived on the tubers, even though they could not be detected by cultivation.

## Conclusions

In this study, none of the used bacterial isolates or isolate mixtures was able to consistently protect potato plants against blackleg disease caused by either *D. solani* or *P. brasiliense*. Still, whole genome sequencing showed that most strains carried genes that enable them to produce secondary metabolites with antibacterial or plant growth promoting properties. *In-vitro* tests confirmed direct antagonism of most isolates against these pathogens. In addition, most isolates could survive on potato tubers from inoculation until the time of planting in the field. Moreover, all strains have been isolated from potato tissue or its vicinity. Therefore, incompatibility with the host is unlikely and it can be assumed that potato tubers have been planted that carried a high density of bacteria with the ability to antagonize *Dickeya* and *Pectobacterium* on the surface. However, inoculation of the antagonists on the outside of the tuber might have been insufficient for biocontrol strains to rapidly colonize the plants through the tuber as most endophytic colonization is known to occur via the rhizosphere. Consequentially the inoculated antagonists were not in direct contact with the pathogen, but were exposed to competition by the indigenous soil microflora.

Instead of relying on inoculated strains for plant protection, the indigenous microflora itself could be used in the protection of potato tubers. Endophytic bacteria are known to mostly originate from the soil and colonize the plant via the roots. Therefore, the soil microbiome can play an important role in disease resistance. Once inside the plant, they can colonize the daughter tubers and might protect the host plant from pathogens in the next field generation..

## Acknowledgements

This research was funded by the Dutch Ministry of Agriculture, Nature and Food Quality (LNV) in the Topsector Program Agriculture & Food (project title: “Effect van de bodem op weerbaarheid van aardappelknollen tegen biotische stress”; project number: AF-17003).

## References

Amir H. and Alabouvette C. (1993). Involvement of soil abiotic factors in the mechanisms of soil suppressiveness to fusarium wilts. Soil Biol Biochem, 25(2), 157–164. 10.1016/0038-0717(93)90022-4.

Andreote F. D., Araújo W. L. d., Azevedo J. L. d., Elsas J. D. v., Rocha U. N. d. and Overbeek L. S. v. (2009). Endophytic colonization of potato *Solanum tuberosum* by a novel competent bacterial endophyte, *Pseudomonas putida* strain P9, and its effect on associated bacterial communities. Appl Environ Microbiol, 75(11), 3396–3406. doi:doi:10.1128/AEM.00491-09.

Andreote F. D., Rocha U. N. d., Araújo W. L., Azevedo J. L. and van Overbeek L. S. (2010). Effect of bacterial inoculation, plant genotype and developmental stage on root-associated and endophytic bacterial communities in potato (*Solanum tuberosum*). Antonie van Leeuwenhoek, 97(4), 389–399. doi:10.1007/s10482-010-9421-9.

Bashan Y., de-Bashan L. E., Prabhu S. R. and Hernandez J.-P. (2014). Advances in plant growth-promoting bacterial inoculant technology: formulations and practical perspectives (1998–2013). Plant Soil, 378(1), 1–33. doi:10.1007/s11104-013-1956-x.

Blin K., et al. (2019). antiSMASH 5.0: updates to the secondary metabolite genome mining pipeline. Nucleic Acids Res, 47(W1), W81–W87. doi:10.1093/nar/gkz310.

Charkowski A. O. (2015). Biology and control of Pectobacterium in potato. Am J Potato Res, 92(2), 223–229. doi:10.1007/s12230-015-9447-7.

Chaumeil P.-A., Mussig A. J., Hugenholtz P. and Parks D. H. (2019). GTDB-Tk: a toolkit to classify genomes with the Genome Taxonomy Database. Bioinformatics, 36(6), 1925–1927. doi:10.1093/bioinformatics/btz848.

Compant S., Duffy B., Nowak J., Clément C. and Barka E. A. (2005). Use of plant growth-promoting bacteria for biocontrol of plant diseases: principles, mechanisms of action, and future prospects. Appl Environ Microbiol, 71(9), 4951–4959.

Czajkowski R., De Boer W., Van Veen J. and Van der Wolf J. (2012). Characterization of bacterial isolates from rotting potato tuber tissue showing antagonism to *Dickeya* sp. biovar 3 *in vitro* and *in planta*. Plant Path, 61(1), 169–182.

Dhar Purkayastha G., Mangar P., Saha A. and Saha D. (2018). Evaluation of the biocontrol efficacy of a *Serratia marcescens* strain indigenous to tea rhizosphere for the management of root rot disease in tea. PLOS ONE, 13(2), e0191761. doi:10.1371/journal.pone.0191761.

Diallo S., Crépin A., Barbey C., Orange N., Burini J.-F. and Latour X. (2011). Mechanisms and recent advances in biological control mediated through the potato rhizosphere. FEMS Microbiol Lett, 75(3), 351–364. 10.1111/j.1574-6941.2010.01023.x.

El-Hassan S. A. and Gowen S. R. (2006). Formulation and delivery of the bacterial antagonist *Bacillus subtilis* for management of lentil vascular wilt caused by *Fusarium oxysporum* f. sp. *lentis*. J Phytopathol, 154(3), 148–155. 10.1111/j.1439-0434.2006.01075.x.

French E., Kaplan I., Iyer-Pascuzzi A., Nakatsu C. H. and Enders L. (2021). Emerging strategies for precision microbiome management in diverse agroecosystems. Nat Plants. doi:10.1038/s41477-020-00830-9.

Hadizadeh I., Peivastegan B., Hannukkala A., van der Wolf J. M., Nissinen R. and Pirhonen M. (2019). Biological control of potato soft rot caused by *Dickeya solani* and the survival of bacterial antagonists under cold storage conditions. Plant Path, 68(2), 297–311. 10.1111/ppa.12956.

Hiddink J. (2022). Bacterieziek stuwt verlagingspercentage pootgoed naar 8.3 procent. Accessed. Krzyzanowska D., et al. (2012). Rhizosphere bacteria as potential biocontrol agents against soft rot caused by various *Pectobacterium* and *Dickeya* spp. strains. J Plant Pathol, 94(2), 367–378.

Krzyzanowska D. M., Maciag T., Siwinska J., Krychowiak M., Jafra S. and Czajkowski R. (2019). Compatible mixture of bacterial antagonists developed to protect potato tubers from soft rot caused by *Pectobacterium* spp. and *Dickeya* spp. Plant Dis, 103(6), 1374–1382. doi:10.1094/pdis-10-18-1866-re.

Lenth R., Singmann H., Love J., Buerkner P. and Herve M. (2018) ’emmeans: estimated marginal means, aka least-square means’ 1.2.3. p.pp. Available at: https://github.com/rvlenth/emmeans (Accessed.

Leong J. (1986). Siderophores: their biochemistry and possible role in the biocontrol of plant pathogens. Annu Rev Phytopathol, 24(1), 187–209.

Mercado-Blanco J. and Bakker P. A. H. M. (2007). Interactions between plants and beneficial *Pseudomonas* spp.: exploiting bacterial traits for crop protection. Antonie van Leeuwenhoek, 92(4), 367–389. doi:10.1007/s10482-007-9167-1.

Oliver J. D. (2010). Recent findings on the viable but nonculturable state in pathogenic bacteria. FEMS Microbiol Rev, 34(4), 415–425. 10.1111/j.1574-6976.2009.00200.x.

Pieterse C. M. J., Zamioudis C., Berendsen R. L., Weller D. M., Van Wees S. C. M. and Bakker P. A. H. M. (2014). Induced systemic resistance by beneficial microbes. Annu Rev Phytopathol, 52(1), 347–375. doi:10.1146/annurev-phyto-082712-102340.

Qiu Z., Egidi E., Liu H., Kaur S. and Singh B. K. (2019). New frontiers in agriculture productivity: Optimised microbial inoculants and *in situ* microbiome engineering. Biotechnol Adv, 37(6), 107371.

Raaijmakers J. M., Vlami M. and de Souza J. T. (2002). Antibiotic production by bacterial biocontrol agents. Antonie van Leeuwenhoek, 81(1), 537. doi:10.1023/a:1020501420831.

Raoul des Essarts Y., et al. (2016). Biocontrol of the potato blackleg and soft rot diseases caused by *Dickeya dianthicola*. Appl Environ Microbiol, 82(1), 268–278. doi:10.1128/aem.02525-15.

RStudio T. (2016) ’RStudio: Integrated Development Environment for R’ 1.4.1717. Boston, MA: RStudio Inc., p.pp. Available at: http://www.rstudio.com/ (Accessed.

Sturz A. V. and Matheson B. G. (1996). Populations of endophytic bacteria which influence host-resistance to Erwinia-induced bacterial soft rot in potato tubers. Plant Soil, 184(2), 265–271. doi:10.1007/bf00010455.

van der Wolf J., Krijger M., Mendes O., Kurm V. and Gros J. (2022). Natural infections of potato plants grown from minitubers with blackleg-causing Soft Rot Pectobacteriaceae. Microorganisms, 10(12), 2504.

van der Wolf J. M., et al. (2017). Virulence of *Pectobacterium carotovorum* subsp. *brasiliense* on potato compared with that of other *Pectobacterium* and *Dickeya* species under climatic conditions prevailing in the Netherlands. Plant Path, 66(4), 571–583. doi:10.1111/ppa.12600.

Van Gijsegem F., van der Wolf J. M. and Toth I. K. (2021) ’Plant diseases caused by Dickeya and Pectobacterium species’. Springer, p.pp. Available at: (Accessed.

Venables W. N. and Ripley B. D. (2013). Modern applied statistics with S-PLUS (trans. New York: Springer Science & Business Media.

Weller D. M. (2007). Pseudomonas biocontrol agents of soilborne pathogens: looking back over 30 years. Phytopathology, 97(2), 250–256.

Zhang W., et al. (2020). Quorum quenching in a novel *Acinetobacter* sp. XN-10 bacterial strain against *Pectobacterium carotovorum* subsp. *carotovorum*. Microorganisms, 8(8), 1100.

