## Supplementary information for "Variable effects of biocontrol bacteria on potato resistance against black leg caused by soft rot Pectobacteriaceae in the field"

a

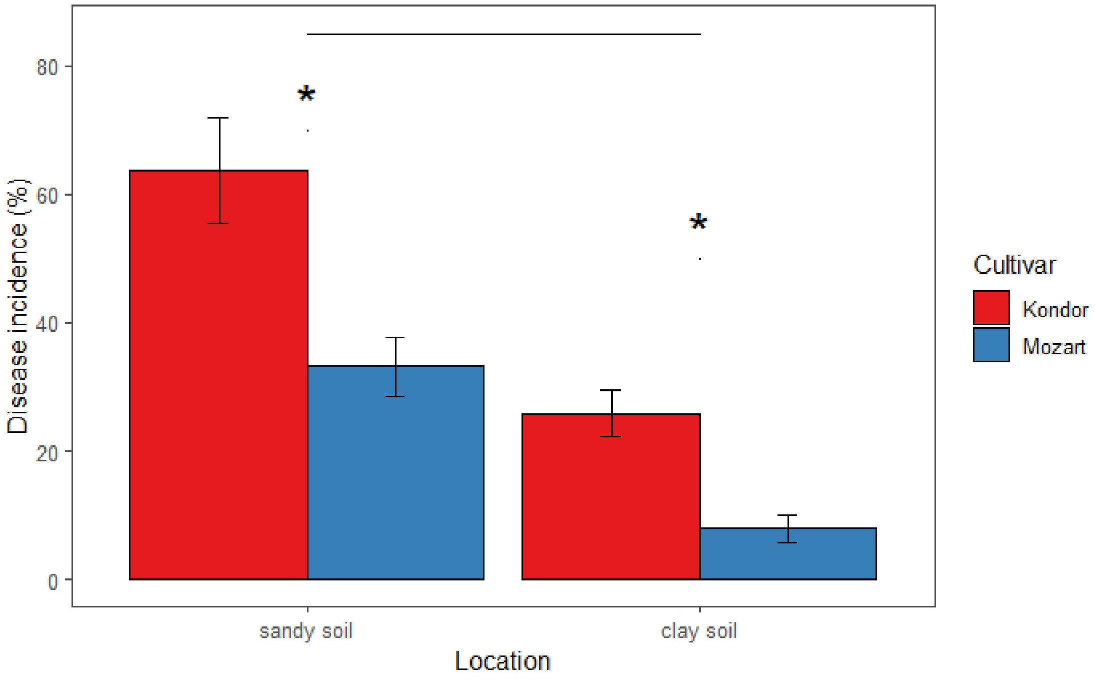

b

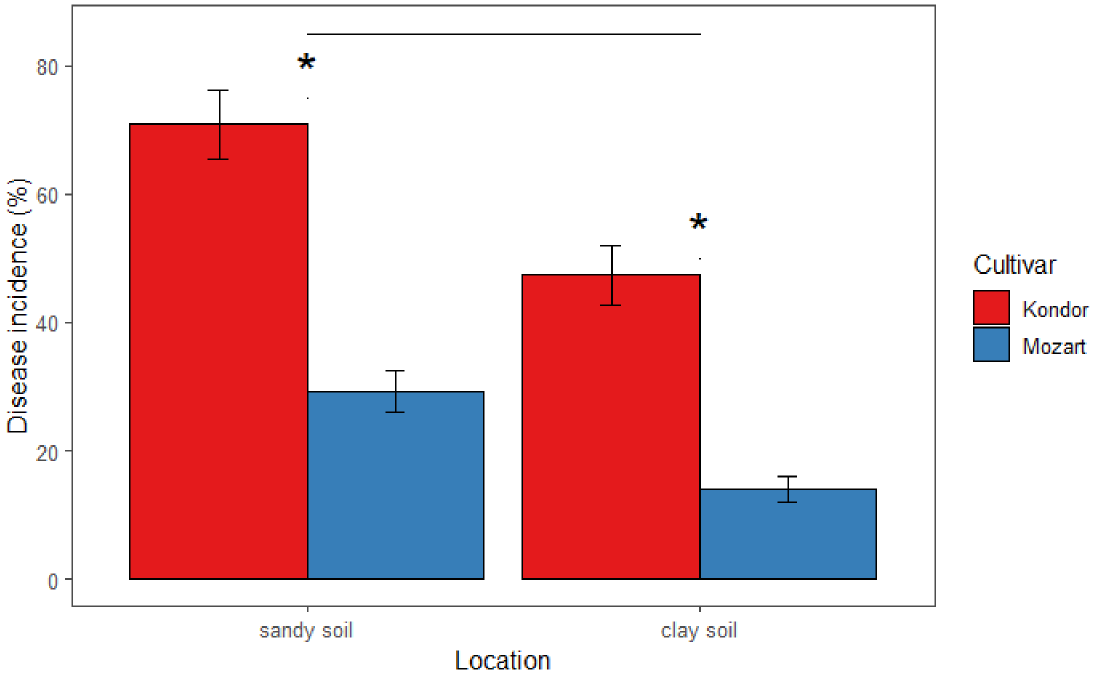

Figure S1: Disease incidence in the year 2020, in both cultivars and locations, averaged over treatment and replicates for a) *D. solani* inoculated tubers and b) *P. brasiliense* inoculated tubers; error bars represent the standard error.

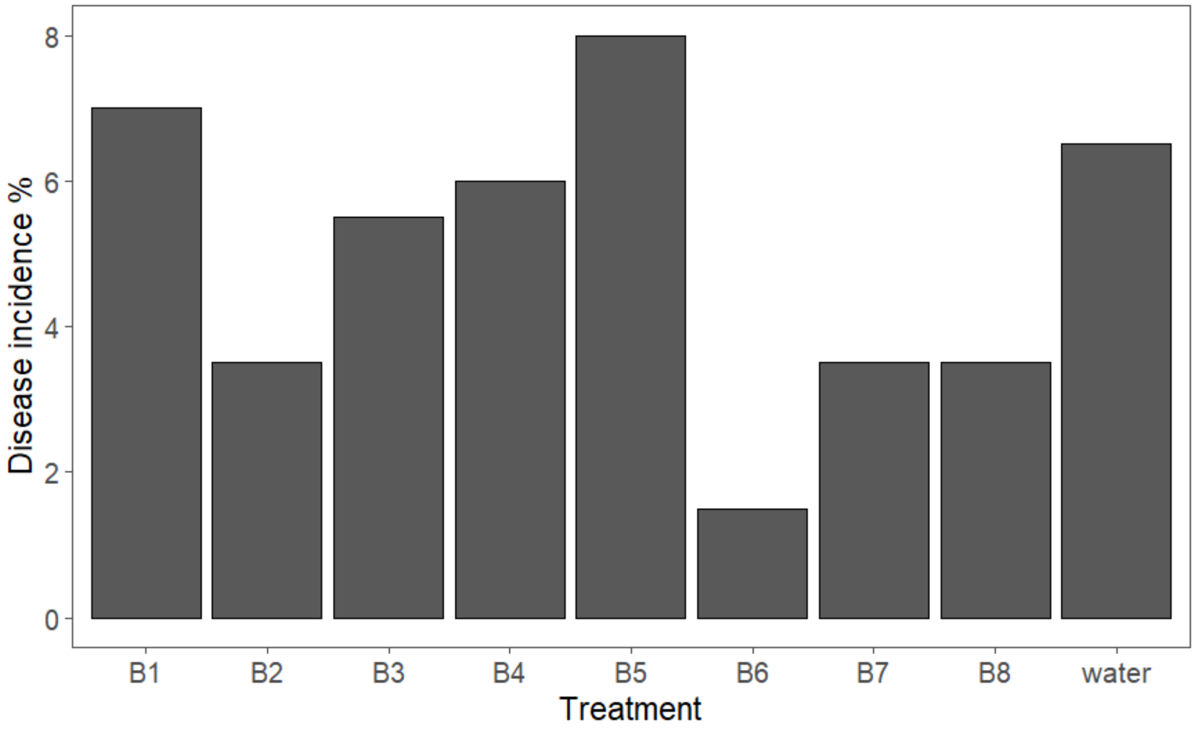

Fig. S2: Disease incidence in 2020, part B, Kondor tubers naturally infected with *P. brasiliense* and treated with 8 different antagonist combinations.

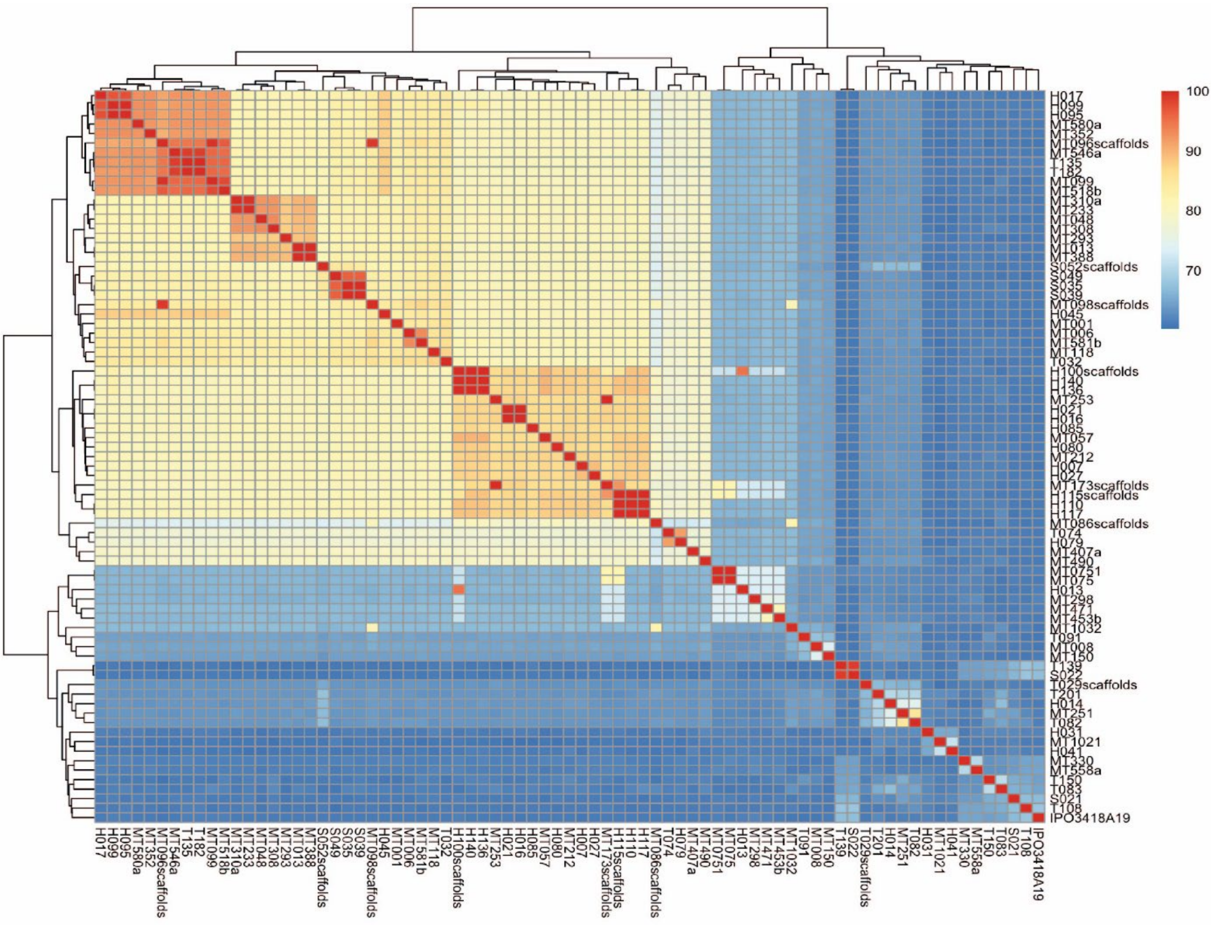

Fig. S3: Average nucleotide analysis of 78 bacterial isolates.

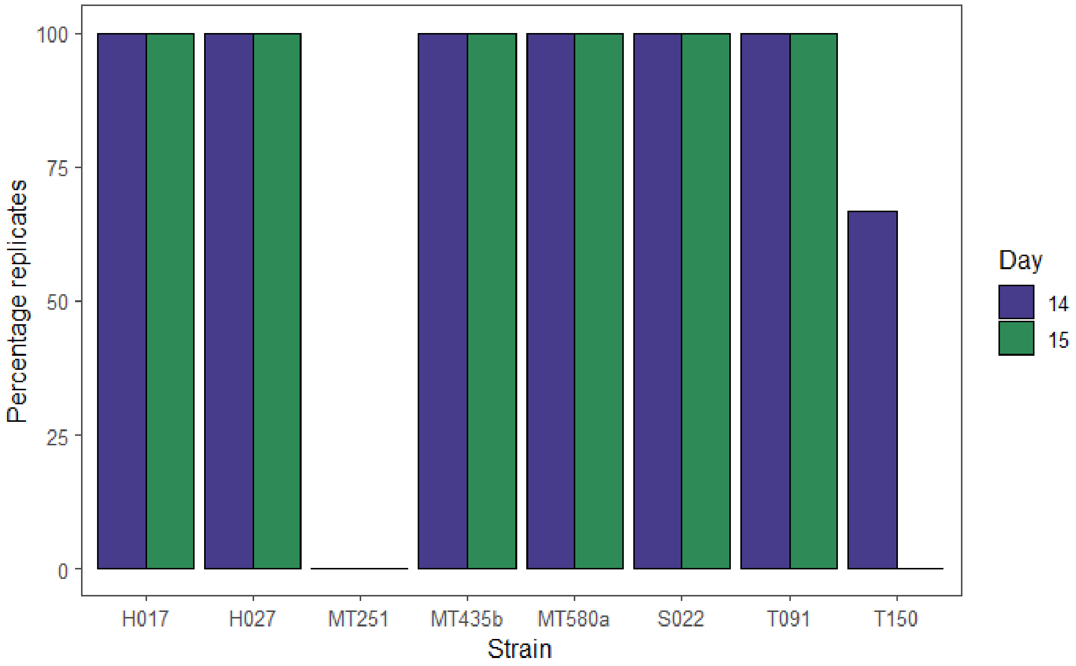

Fig S4: Percentage of replicates (n=3) in which isolates were identified after 14 and 15 days after inoculation on tubers as single strains.

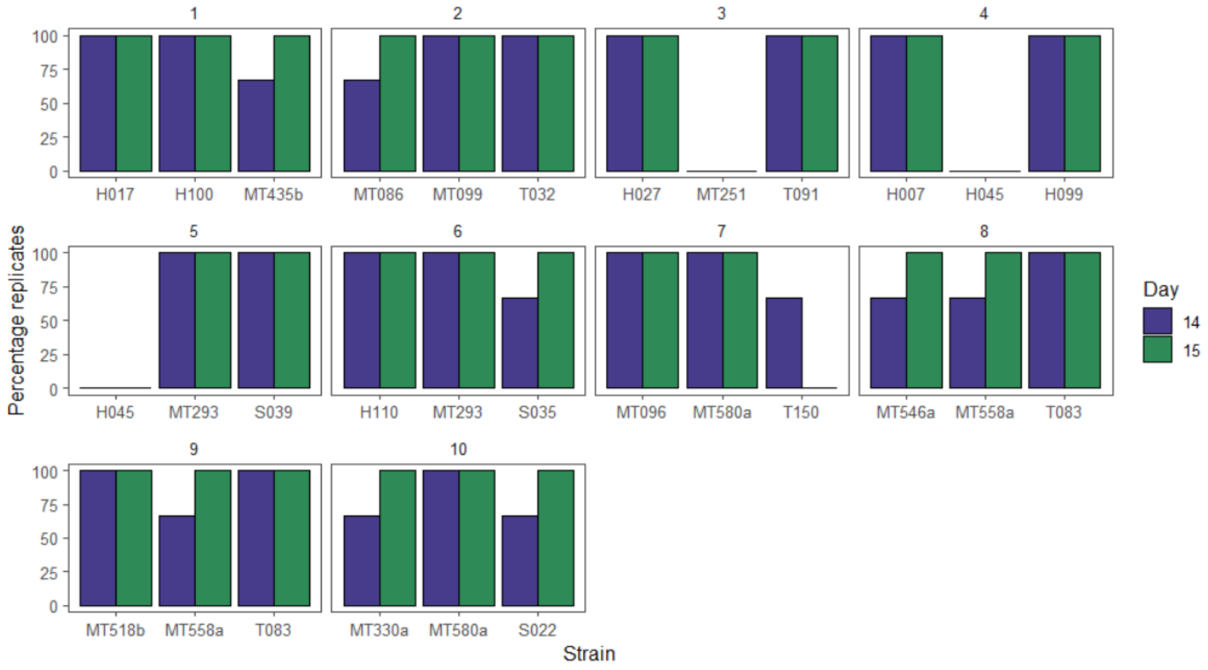

Fig S5: Percentage of replicates (n=3) in which isolates were identified after 14 and 15 days after inoculation on tubers in mixes of three.

Table S5: Average disease incidence in 2020, in the two soil types and cultivars, averaged over treatment and replicates and in cultivar Kondor. Statistical results of an Anova.

| Year | Part | Pathogen | Cultivar | Location | Average disease incidence (%) | se | contrast | Chi-sq | p |
|---|---|---|---|---|---|---|---|---|---|
| 2020 | A | *P. brasiliense* | | sandy soil | 50.07 | 3.99 | sandy-clay soil | 98.69 | <0.01 |
| 2020 | A | *P. brasiliense* | | clay soil | 30.68 | 3.18 | | | |
| 2020 | A | *P. brasiliense* | Kondor | | 59.11 | 3.77 | Kondor-Mozart | 357.87 | <0.01 |
| 2020 | A | *P. brasiliense* | Mozart | | 21.65 | 2.14 | | | |
| 2020 | A | *D. solani* | | sandy soil | 49.19 | 5.33 | sandy-clay soil | 159.69 | <0.01 |
| 2020 | A | *D. solani* | | clay soil | 17.38 | 2.6 | | | |
| 2020 | A | *D. solani* | Kondor | | 44.79 | 5.31 | Kondor-Mozart | 97.89 | <0.01 |

| | | | | | | | | | |
|---|---|---|---|---|---|---|---|---|---|
| 2020 | A | *D. solani* | Mozart | | 20.57 | 3.25 | | | |
| 2020 | C | *P. brasiliense* | | sandy soil | 55.39 | 2.62 | sandy-clay soil | 271.75 | <0.01 |
| 2020 | C | *P. brasiliense* | | clay soil | 35.04 | 1.94 | | | |
| 2020 | C | *P. brasiliense* | Kondor | | 61.35 | 2.43 | Kondor-Mozart | 659.37 | <0.01 |
| 2020 | C | *P. brasiliense* | Mozart | | 28.21 | 1.54 | | | |
| 2020 | C | *D. solani* | | sandy soil | 55.91 | 2.41 | sandy-clay soil | 297.61 | <0.01 |
| 2020 | C | *D. solani* | | clay soil | 34.56 | 1.89 | | | |
| 2020 | C | *D. solani* | Kondor | | 62.61 | 2.03 | Kondor-Mozart | 736.73 | <0.01 |
| 2020 | C | *D. solani* | Mozart | | 27.64 | 1.68 | | | |

Table S6: Average disease incidence in 2020, in tubers inoculated with *P. brasiliense* and with treatments that significantly decreased disease incidence compared to the positive control in sandy soil averaged over cultivar and replicates and in cultivar Kondor averaged over locations and replicates. Statistical results of a pairwise comparison with the positive control.

| **Location** | **Pathogen** | **Treatment** | **Average disease incidence** | **se** | **z-ratio** | **p-value** |
|---|---|---|---|---|---|---|
| sandy soil | *P. brasiliense* | 130 | 16.96 | 4.46 | -3.59 | 0.02 |
| sandy soil | *P. brasiliense* | 127 | 18.75 | 6.25 | -3.55 | 0.02 |
| sandy soil | *P. brasiliense* | 68 | 12.92 | 0.42 | -3.85 | 0.01 |
| sandy soil | *P. brasiliense* | 37 | 18.75 | 6.25 | -3.55 | 0.02 |
| sandy soil | *P. brasiliense* | Pos. CTRL | 56.87 | 8.89 | | |
| **Cultivar** | ***Pathogen*** | **Treatment** | **Disease incidence** | **se** | **z-ratio** | **p-value** |
| Kondor | *P. brasiliense* | 131 | 29.38 | 10.63 | -3.54 | 0.02 |
| Kondor | *P. brasiliense* | 130 | 27.88 | 2.88 | -4.23 | <0.01 |
| Kondor | *P. brasiliense* | 129 | 36.65 | 17.90 | -3.68 | 0.01 |

| | | | | | | |
|---|---|---|---|---|---|---|
| Kondor | *P. brasiliense* | 127 | 37.5 | 6.25 | -3.54 | 0.02 |
| Kondor | *P. brasiliense* | 69 | 6.67 | 6.67 | -3.81 | 0.01 |
| Kondor | *P. brasiliense* | 68 | 12.5 | 6.25 | -4.53 | <0.01 |
| Kondor | *P. brasiliense* | 35 | 16.67 | 16.67 | -3.81 | 0.01 |
| Kondor | *P. brasiliense* | 34 | 12.5 | 12.5 | -4.13 | <0.01 |
| Kondor | *P. brasiliense* | Pos. CTRL | 40.96 | 9.30 | | |

Table S7: Average disease incidence in 2021, in cultivar Mozart and Kondor and in sandy soil and clay soil, averaged over treatments and replicates. Statistical results of a pairwise comparison with the positive control.

| **Year** | **Part** | **Pathogen** | **Cultivar** | **Location** | **Average disease incidence (%)** | **se** | **contrast** | **z-ratio** | **p** |
|---|---|---|---|---|---|---|---|---|---|
| 2021 | A | *P. brasiliense/D. solani* | Kondor | sandy soil | 65.32 | 5.23 | sandy-clay soil | -0.16 | 0.88 |
| 2021 | A | *P. brasiliense/D. solani* | Kondor | clay soil | 65.84 | 6.26 | | | |
| 2021 | A | *P. brasiliense/D. solani* | Mozart | sandy soil | 33.65 | 3.89 | sandy-clay soil | 6.03 | <0.01 |
| 2021 | A | *P. brasiliense/D. solani* | Mozart | clay soil | 15.68 | 1.85 | | | |
| 2021 | B1 | *P. brasiliense/D. solani* | Kondor | sandy soil | 51.24 | 6.53 | sandy-clay soil | 0.36 | 0.72 |
| 2021 | B1 | *P. brasiliense/D. solani* | Kondor | clay soil | 49.79 | 7.13 | | | |
| 2021 | B1 | *P. brasiliense/D. solani* | Mozart | sandy soil | 30.84 | 4.39 | sandy-clay soil | 3.74 | <0.01 |
| 2021 | B1 | *P. brasiliense/D. solani* | Mozart | clay soil | 17.17 | 2.62 | | | |

| | | | | | | | | | |
|---|---|---|---|---|---|---|---|---|---|
| 2021 | B2 | *P. brasiliense/D. solani* | Kondor | sandy soil | 44.38 | 6.49 | sandy-clay soil | 0.6 | 0.55 |
| 2021 | B2 | *P. brasiliense/D. solani* | Kondor | clay soil | 41.98 | 4.86 | | | |
| 2021 | B2 | *P. brasiliense/D. solani* | Mozart | sandy soil | 34.96 | 4.86 | sandy-clay soil | 4.85 | <0.01 |
| 2021 | B2 | *P. brasiliense/D. solani* | Mozart | clay soil | 17.3 | 2.77 | | | |
| 2021 | C | *P. brasiliense* | Kondor | sandy soil | 73.46 | 2.45 | sandy-clay soil | -5.03 | <0.01 |
| 2021 | C | *P. brasiliense* | Kondor | clay soil | 86.03 | 3.2 | | | |
| 2021 | C | *P. brasiliense* | Mozart | sandy soil | 32.97 | 1.79 | sandy-clay soil | 3.67 | <0.01 |
| 2021 | C | *P. brasiliense* | Mozart | clay soil | 22.46 | 1.47 | | | |
| 2021 | C | *D. solani* | Kondor | sandy soil | 70.17 | 2.09 | sandy-clay soil | 8.83 | <0.01 |
| 2021 | C | *D. solani* | Kondor | clay soil | 42.99 | 2.57 | | | |
| 2021 | C | *D. solani* | Mozart | sandy soil | 38.31 | 4.35 | sandy-clay soil | 0.01 | 0.99 |
| 2021 | C | *D. solani* | Mozart | clay soil | 15.17 | 2.08 | | | |
| 2021 | D | *P. brasiliense* | Kondor | sandy soil | 63.05 | 3.25 | sandy-clay soil | -3.88 | <0.01 |
| 2021 | D | *P. brasiliense* | Kondor | clay soil | 71.8 | 3.61 | | | |
| 2021 | D | *P. brasiliense* | Mozart | sandy soil | 36.1 | 2.4 | sandy-clay soil | 7.32 | <0.01 |
| 2021 | D | *P. brasiliense* | Mozart | clay soil | 19.77 | 1.47 | | | |
| 2021 | D | *D. solani* | Kondor | sandy soil | 23.23 | 3.05 | sandy-clay soil | 10.01 | <0.01 |
| 2021 | D | *D. solani* | Kondor | clay soil | 38.89 | 2.16 | | | |
| 2021 | D | *D. solani* | Mozart | sandy soil | 46.34 | 2.77 | sandy-clay soil | 11.75 | <0.01 |

| | | | | | | |
|---|---|---|---|---|---|---|
| 2021 | D | *D. solani* | Mozart | clay soil | 18.55 | 1.49 |

Table S8: BUSCO analysis results for the whole genomes of the 25 selected antagonist strains and identification by GTDB-Tk

| **Strain ID** | **ID** | **Order** | **Complete** | **Fragmented** | **Missing** | **Duplicated** | **Identity** |
|---|---|---|---|---|---|---|---|
| A5 | H007 | Pseudomonadales | 99.7 | 0 | 0.2 | 0.1 | *Pseudomonas fluorescens* |
| A1 | H017 | Pseudomonadales | 99.6 | 0.3 | 0.1 | 0.1 | *Pseudomonas koreensis* |
| C5 | H027 | Pseudomonadales | 99.8 | 0.1 | 0.1 | 0.1 | *Pseudomonas sp.* |
| A6 | H045 | Pseudomonadales | 99.7 | 0 | 0 | 0.3 | *Pseudomonas putida* |
| A3 | H099 | Pseudomonadales | 90.2 | 3.8 | 6 | 2.7 | *Pseudomonas koreensis* |
| A11 | H100 | Pseudomonadales | 100 | 0 | 0 | 7.9 | *Pseudomonas marginalis+ Pantoea vagans* |
| B1 | H110 | Pseudomonadales | 99.4 | 0.3 | 0.2 | 0.1 | *Pseudomonas sp.* |
| C1 | MT086 | Xanthomonadales | 99.4 | 0.1 | 0.5 | 1.4 | *Stenotrophomonas sp.* |
| N8 | MT096 | Pseudomonadales | 98.4 | 0.9 | 0.7 | 91.9 | *Pseudomonas sp.* |
| C2 | MT099 | Pseudomonadales | 99.7 | 0.1 | 0.2 | 0.1 | *Pseudomonas sp.* |
| C18 | MT251 | Micrococcales | 99.2 | 0.2 | 0.6 | 0.7 | *Curtobacterium sp.* |
| C20 | MT293 | Pseudomonadales | 99.8 | 0.1 | 0.1 | 0.1 | *Pseudomonas sp.* |
| N1 | MT330a | Bacillales | 98.4 | 1.3 | 0.3 | 0.4 | *Paenibacillus sp.* |
| A21 | MT453b | Enterobacterales | 99.7 | 0.2 | 0.1 | 0.2 | *Serratia fonticola* |
| N5 | MT518b | Pseudomonadales | 99.8 | 0.1 | 0.1 | 0.1 | *Pseudomonas sp.* |
| N4 | MT546a | Pseudomonadales | 99.7 | 0.3 | 0 | 0.1 | *Pseudomonas sp.* |
| N2 | MT558a | Bacillales | 98.9 | 0.7 | 0.4 | 1.3 | *Paenibacillus polymyxa* |
| N12 | MT580a | Pseudomonadales | 99.8 | 0.1 | 0.1 | 0.1 | *Pseudomonas sp.* |

| | | | | | | | |
|---|---|---|---|---|---|---|---|
| N3 | S022 | Bacillales | 99.8 | 0 | 0.2 | 0.2 | *Bacillus altitudinis* |
| B7 | S035 | Pseudomona dales | 99.7 | 0.1 | 0.2 | 0.1 | *Pseudomonas fluorescens* |
| C16 | S039 | Pseudomona dales | 99.7 | 0.1 | 0.2 | 0.1 | *Pseudomonas fluorescens* |
| C3 | T032 | Pseudomona dales | 100 | 0 | 0 | 0 | *Pseudomonas sp.* |
| N11 | T083 | Staphylococ cales | 99.8 | 0 | 0.2 | 0 | *Staphylococcus vitulinus* |
| C15 | T091 | Sphingomon adales | 99.9 | 0.1 | 0 | 0.6 | *Sphingobium yanoikuyae* |
| N10 | T150 | Staphylococ cales | 99.7 | 0 | 0.3 | 0.4 | *Staphylococcus sp.* |
